# Cell Type–Dependent Uptake of Extracellular Vesicles Independent of Cellular Origin

**DOI:** 10.64898/2026.05.19.726167

**Authors:** Doste R. Mamand

## Abstract

Extracellular vesicles (EVs) are promising nanocarriers for therapeutic delivery; however, the factors governing EV uptake by recipient cells remain incompletely understood. In this study, we investigated whether EV internalization is primarily influenced by donor-cell origin or recipient-cell phenotype. Fluorescently labeled EVs derived from HEK293T, or SKBR-3 cells were incubated with a range of human epithelial, immune, and murine cancer cell lines at different doses and time points. HEK293T-derived EVs showed highly variable uptake across recipient cells, with hepatocellular carcinoma cell lines Huh7 and HepG2 exhibiting the highest internalization, while parental HEK293T cells showed the lowest. THP-1 immune cells also demonstrated strong uptake, whereas Jurkat cells showed moderate uptake. In murine melanoma models, Yummer cells internalized more EVs than B16F10 cells. Importantly, similar uptake trends were observed using SKBR-3-derived EVs, where Huh7 and HepG2 again displayed the highest uptake despite originating from a different donor cell source. EV internalization increased with dose and incubation time until saturation at higher concentrations. Together, these results demonstrate that EV uptake is predominantly determined by recipient-cell characteristics rather than EV source. These findings provide important mechanistic insight for the development of EV-based therapeutics and suggest that optimizing recipient-cell targeting is essential for efficient vesicle-mediated delivery.

**Graphical abstract:** EV uptake is determined by cell membrane properties rather than by the source of the EVs. The image was created by Biorender.

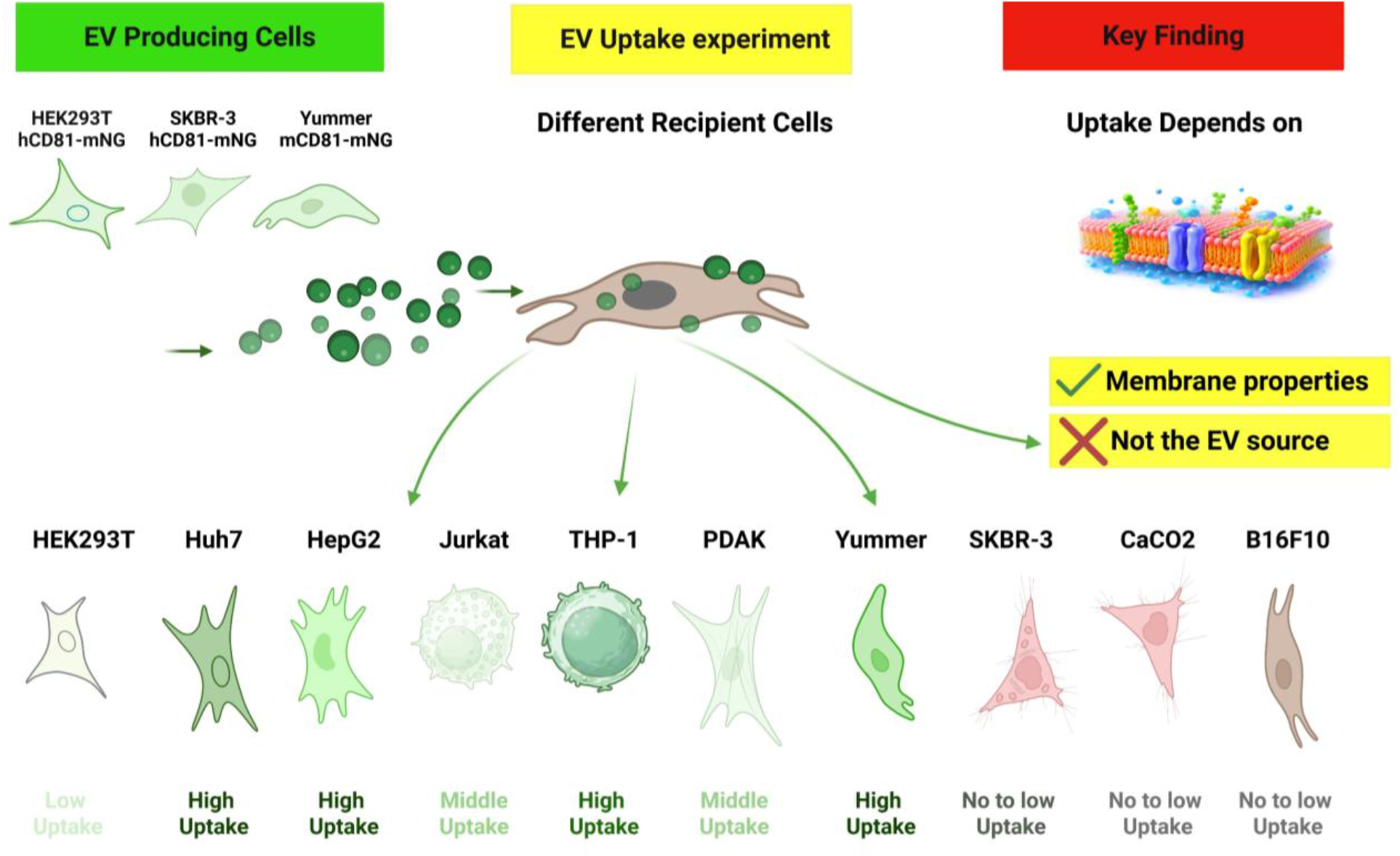

## 1. Introduction

Cellular uptake of nanoparticles, including EVs, is a fundamental biological process that underpins intercellular communication and the transfer of molecular information between cells[1], [2]. EVs are lipid bilayer-enclosed particles naturally released by cells and are known to carry proteins, lipids, and nucleic acids that can modulate the behaviour and function of recipient cells[3], [4]. Due to these properties, EVs have emerged as promising candidates for applications in diagnostics, drug delivery, and regenerative medicine[5], [6]. The mechanisms governing EV uptake are complex and multifactorial, involving a range of processes such as clathrin-mediated endocytosis, caveolin-dependent uptake, macropinocytosis, phagocytosis, and direct membrane fusion. These pathways are influenced by both the biochemical composition of EVs and the physiological characteristics of recipient cells[7], [8]. Historically, considerable emphasis has been placed on the role of EV origin specifically, the idea that vesicles derived from certain cell types possess inherent targeting capabilities that dictate their uptake specificity[9], [10], [11]. However, increasing evidence suggests that this perspective may be incomplete. Emerging studies indicate that recipient cell properties such as membrane receptor expression, lipid composition, cytoskeletal organization, and metabolic activity—may play a dominant role in determining EV uptake efficiency. This implies that EV internalization may be less dependent on where the vesicles originate and more dependent on how receptive the target cells are to vesicle interaction and internalization[12], [13], [14], [15]. This shift in understanding has significant implications for both basic biology and translational applications. If EV uptake is primarily governed by recipient cell type, strategies for EV-based therapeutics may need to focus more on modifying or selecting target cells rather than engineering vesicle origin alone[16], [17], [18], [19]. Furthermore, this insight could help explain inconsistencies observed in previous studies where EV uptake did not correlate strongly with vesicle source [20], [21], [22], [23]. Despite these advances, a systematic and controlled comparison of EV uptake across multiple donor and recipient cell combinations remains lacking. Many existing studies are limited by variability in experimental conditions, EV isolation methods, or insufficient cross-comparisons between cell types. Therefore, there is a critical need for a well-controlled investigation that directly addresses whether EV uptake is predominantly dictated by recipient cell identity

In this project, we aim to fill this gap by conducting a comprehensive analysis of EV uptake across diverse cellular systems. By standardizing EV preparation, labelling, and exposure conditions, and by employing robust quantitative and flow Cytometry-based methods, we seek to isolate the key variables influencing uptake. Ultimately, this study will provide deeper insight into the fundamental principles governing EV–cell interactions. The findings are expected to contribute to the refinement of EV-based delivery strategies, enhance our understanding of intercellular communication, and inform the development of more effective and targeted therapeutic approaches.

## 2. Materials and Methods

### 2.1 Materials

Dulbecco’s Modified Eagle Medium (DMEM, high glucose; Catalog No. 11965092), Roswell Park Memorial Institute medium (RPMI-1640; Catalog No. 11875093), fetal bovine serum (FBS), serum-free Opti-MEM (Catalog No. 31985070), and phosphate-buffered saline (PBS; Catalog No. 10010015) were obtained from Gibco (USA). A 1% antibiotic–antimycotic (anti-anti) solution (Catalog No. 15240062) was purchased from Gibco (USA). Mycoplasma detection kits were sourced from Eurofins and routinely used to screen cell cultures for contamination.

### 2.2 Cell Culture

Human embryonic kidney cells (HEK293T), SKBR-3 human breast cancer cells, pancreatic ductal adenocarcinoma (PDAK) cells, HepG2 human hepatic cancer cells, Huh7 human hepatocellular carcinoma cells, Caco-2 human colorectal adenocarcinoma cells, and the murine melanoma cell lines B16F10 and Yummer (all obtained from ATCC) were cultured in DMEM supplemented with 10% FBS and 1% antibiotic–antimycotic solution. Cells were maintained in 150 mm tissue culture dishes (standard surface; Sarstedt, Cat. No. 83.3903.300) at 37 °C in a humidified incubator with 5% CO_2_ for 24 h to permit cell attachment and recovery. Subsequently, the culture medium was replaced with serum-free Opti-MEM containing 1% antibiotic–antimycotic solution to enable extracellular vesicle (EV) production under serum-free conditions. The conditioned medium (CM) was harvested after 48 h, following which it was purified and concentrated for subsequent use.

### 2.3 Plasmid generation

Plasmids encoding human CD81 or mouse CD81 fused to mNeonGreen (mNG) were generated using the lentiviral p2CL9IPw5 backbone vector. The amino acid sequences for human CD81, mouse CD81, and mNG were verified using UniProt. Synthesized gene fragments (Integrated DNA Technologies) were cloned downstream of the spleen focus-forming virus (SFFV) promoter within the p2CL9IPw5 vector using standard restriction enzyme cloning strategies[24]. The constructs additionally contained an internal ribosomal entry site (IRES)-puromycin resistance cassette for stable cell selection. All plasmid constructs were verified by Sanger sequencing performed by Eurofins Genomics prior to use in lentiviral production.

### 2.4 Generating HEK293T, SKBR-3, and Yummer cells stably expressing CD81-mNG EVs

Lentiviral supernatants were generated as previously described in reference [24]. Briefly, HEK293T cells were co-transfected with the p2CL9IPw5 construct encoding human or mouse CD81 tagged with mNeonGreen (mNG), the pCD/NL-BH helper construct, and the pcoPE human foamy virus envelope construct using polyethyleneimine (PEI). At 18 h post-transfection, expression from the human cytomegalovirus immediate-early enhancer/promoter was enhanced by treating the cells with 10 mM sodium butyrate (Sigma-Aldrich) for 7 h. The medium was then replaced, and the viral supernatant was collected 22 h later. Viral particles were concentrated by centrifugation at 25,000 × g for 90 min at 4°C. After removing the supernatant, the viral pellet was resuspended in 1 mL Iscove’s Modified Dulbecco’s Medium supplemented with 20% FBS and 1% antibiotic–antimycotic solution. Virus stocks were stored at −80°C until use. Cells were then transduced with the lentivirus and cultured under puromycin selection for at least five passages to generate stable cell lines.

### 2.5 Isolation of EVs

HEK293T, SKBR-3, and Yummer cells were seeded at a density of 1 × 10^7 cells in 20 mL DMEM per 150 × 20 mm Petri dish (SARSTEDT AG & Co. KG, Germany) and cultured under the conditions described previously[25]. After 24 h, the culture medium was replaced with serum-free Opti-MEM. Following an additional 48 h incubation period, the conditioned media (CM) were collected for subsequent EV isolation and purification as described below.

### 2.6 Purification of EVs

EVs were isolated from conditioned media (CM) using sequential centrifugation, filtration, ultrafiltration, and size exclusion chromatography (SEC). Initially, the CM was pre-cleared by centrifugation at 700 × g for 5 min, followed by centrifugation at 2,000 × g for 10 min to remove cells, debris, and larger particles. Unless otherwise specified, samples were subsequently filtered through a 0.22 µm cellulose acetate bottle-top filter (SARSTEDT AG & Co. KG, Germany) to eliminate remaining large particles.

The filtered CM was concentrated by tangential flow filtration using a MicroKross ultrafiltration module (20 cm^2^, 300 kDa molecular weight cut-off; Spectrum Laboratories). The resulting retentate was further concentrated using Amicon Ultra-15 centrifugal filter units with a 10 kDa molecular weight cut-off (MilliporeSigma) to a final volume of 100 µL, as previously described by Mamand et al.[26]. Subsequently, EVs were purified by size exclusion chromatography using qEV 70 nm columns for EV isolation (Izon Science). Fractions 4 and 5 were collected in a total volume of 2 mL and further concentrated using Amicon® Ultra-15 centrifugal filter units (10 kDa; Merck). Purified EV preparations were stored at −80°C until further analysis.

### 2.7 Nanosight tracking Analysis

Nanoparticle tracking analysis (NTA) measurements were performed using the Particle Metrix ZetaView system as performed before[25], [27]. Samples were diluted in 0.22 µm-filtered PBS to a final volume of 1 mL. Optimal measurement concentrations were established through preliminary testing to obtain particle counts ranging from 140 to 200 particles per frame. The manufacturer’s recommended default software settings for EV analysis were applied. These settings included a camera sensitivity of 80 and a shutter speed of 100. For each sample, two measurement cycles were carried out across 11 cell positions, with 60 frames recorded per position using the high video setting. Measurement parameters were configured as follows: autofocus enabled, camera sensitivity set to 80.0, shutter speed maintained at 100, scattering intensity adjusted to 4.0, and cell temperature maintained at 24°C. Following video acquisition, the recorded data were processed and analyzed using the integrated ZetaView Software version 8.02.31.

### 2.8 Electron microscopy of EV morphology

Three microliters of sample were applied to glow-discharged, carbon-coated, Formvar-stabilized 400-mesh copper grids (easiGlow™, Ted Pella) and incubated for approximately 30 s [25]. Excess sample was blotted off, followed by a Milli-Q water wash. The grids were negatively stained with 1% ammonium molybdate and imaged using an HT7800-Xarosa transmission electron microscope (TEM; Hitachi High-Technologies) operated at 80 kV and equipped with a 4 MP Veleta CCD camera (Olympus Soft Imaging Solutions GmbH).

### 2.9 Imaging flow cytometry for characterization of EV surface markers

EV surface marker characterization was performed using imaging flow cytometry as previously described[26]. Purified EVs were diluted in Dulbecco’s phosphate-buffered saline (DPBS) to a concentration of 1 × 10^10 particles/mL. A total of 2.5 × 10^8 EVs in 25 µL were incubated with 8 nM allophycocyanin (APC)-conjugated antibodies against the EV-associated surface marker CD81 (Miltenyi Biotec). Samples were incubated overnight at room temperature in the dark in sealed 96-well plates (SARSTEDT AG & Co. KG, Germany). Following incubation, the samples were diluted in 1% human albumin and trehalose buffer (PBS-HAT) as previously described (reference 3) to achieve a final concentration of 1 × 10^7 EVs/mL in a total volume of 100 µL without further antibody addition. Single-vesicle imaging and small-particle analysis were carried out using the Amnis CellStream imaging flow cytometer from Luminex Corporation. Data acquisition and analysis were performed using FlowJo v10.8 software (FlowJo LLC).

### 2.10 Western blotting for EV markers

Western blotting was performed to evaluate the presence of the intravesicular protein TSG101 in EVs derived from HEK293T and SKBR-3 cells[18]. Briefly, 2 × 10^10 EVs (equivalent to approximately 15 µg EV protein) and 2 × 10^6 HEK293T or SKBR-3 cells were lysed in 100 µL radioimmunoprecipitation assay (RIPA) buffer (Bio-Rad Laboratories, Hercules, CA, USA). Lysates were incubated on ice for 30 min with intermittent vortexing for 10 s every 5 min. The EV and cell lysates were mixed with 8 µL loading buffer composed of 10% glycerol, 8% sodium dodecyl sulfate (SDS), 0.5 M dithiothreitol (DTT), and 0.4 M sodium carbonate. Samples were heated at 70°C for 5 min prior to separation on NuPAGE™ 4–12% Bis-Tris gels (Invitrogen) and electrophoresed at 120 V for 1.5 h. Proteins were subsequently transferred to iBlot transfer membranes using the iBlot 2 Dry Blotting System for 7 min. Membranes were blocked for 1 h at room temperature using Odyssey Blocking Buffer (LI-COR Biosciences, Lincoln, NE, USA) and incubated overnight at 4°C with freshly prepared anti-TSG101 primary antibody (MA1-23296, Invitrogen) diluted 1:500. Following four washes with 0.1% TBS-T for 5 min each under agitation, membranes were incubated for 1 h with a 1:10,000 diluted secondary antibody (goat anti-human, H10007). After three PBS washes, protein bands were visualized using the LI-COR Odyssey CLx Imaging System.

### 2.11 Uptake experiment

For each uptake experiment, 15,000 cells from each cell line were seeded per well in 96-well plates (Catalog No. 83.3924.005, Sarstedt). Adherent cells were cultured in complete DMEM, whereas suspension cells were maintained in complete RPMI medium. After 24 h of incubation, EVs were added at concentrations of 1 × 10^8, 1 × 10^9, and 1 × 10^10 EVs per well. Cells were incubated with EVs for 2 h or 4 h at 37°C under standard cell culture conditions. Following incubation, cells were washed twice with PBS to remove unbound EVs. Adherent cells were detached by trypsinization for 5 min, while suspension cells were collected directly after washing. Cells were subsequently resuspended in fresh culture medium and analyzed by flow cytometry using a MACSQuant 16 flow cytometer (Miltenyi Biotec). At least 5000 to 10,000 events were recorded per sample. Forward and side scatter parameters were used to gate viable cell populations and exclude debris, and fluorescence intensity was measured to evaluate EV uptake by recipient cells. Data acquisition and analysis were performed using FlowJo v10.8 software (FlowJo LLC).

### 2.12 Statistical Analysis

Data are presented as mean ± standard deviation (SD) from three biological replicates. No data were excluded from the analyses. Statistical analyses were performed using one-way analysis of variance (ANOVA) followed by Tukey’s and Dunnett’s multiple-comparisons tests, and statistical significance was defined as p < 0.05. All analyses were conducted using GraphPad Prism version 10.

## 3. Results

To investigate the uptake of EVs between cell lines in vitro, three donor cell lines were transduced with a lentivirus encoding CD81-mNeonGreen (mNG). This generated stable cell lines expressing fluorescently labeled CD81-mNG EVs (Figure 1, Step 1). The engineered donor cells were subsequently cultured in EV-free serum media to enable secretion of mNG-labeled EVs into the conditioned medium. The released EVs were then isolated and purified by SEC as illustrated in Figure 1 (Step 2). Following isolation, the purified EVs were characterized to confirm their quality, size distribution, and marker expression, as shown in Figure 1 (Step 3). Next, ten different recipient cancer cell lines were utilized for the uptake study, including HEK293T, SKBR-3, Yummer, B16F10, HepG2, Huh7, Caco-2, THP-1, Jurkat, and PDAK, as indicated in Figure 1 (Step 4). In the final step, EV uptake by recipient cells was evaluated using fluorescence-based analysis to determine differences in cellular internalization efficiency among the tested cell lines.

**Figure 1.**
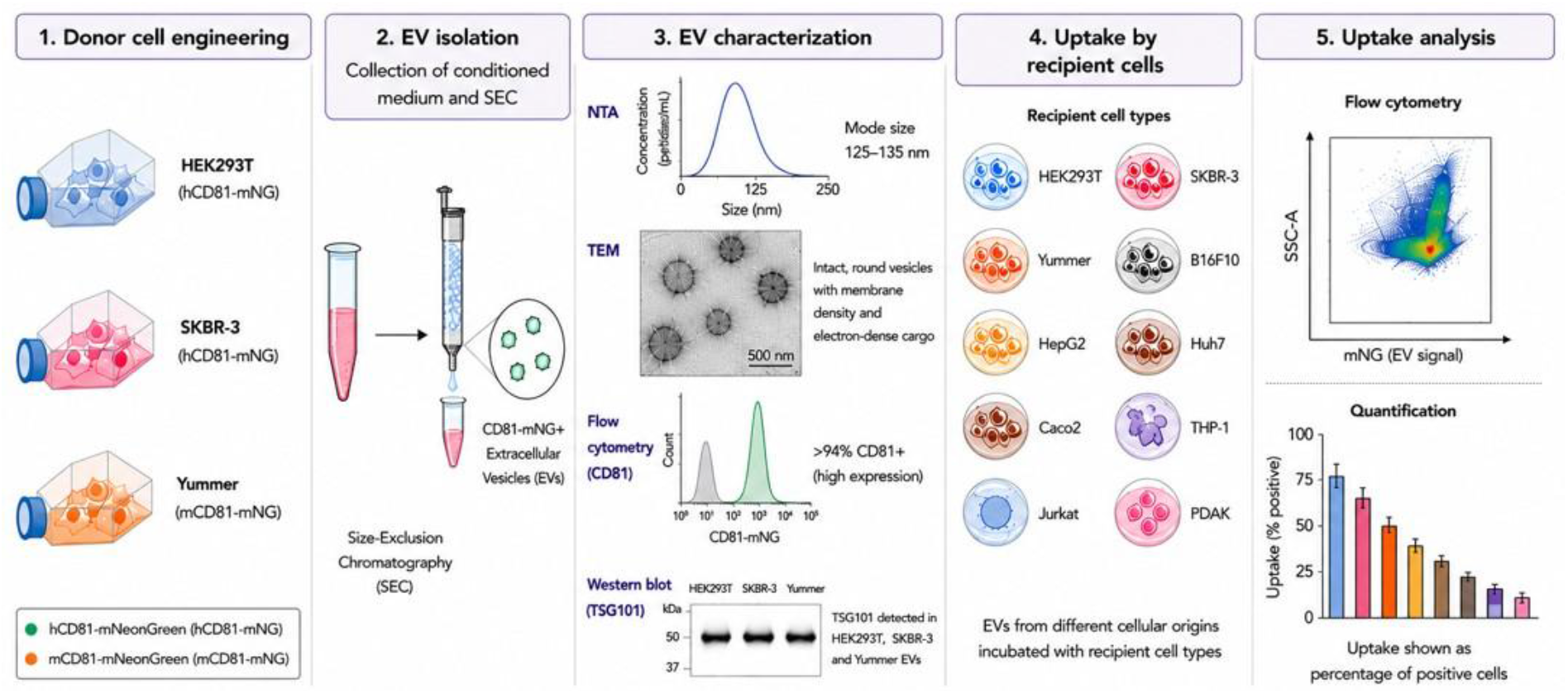
Experimental workflow for EV generation, isolation, characterization, and uptake analysis. Schematic overview of the experimental design used to investigate EV uptake across multiple recipient cell types. Donor cell lines (HEK293T, SKBR-3, and Yummer) were genetically engineered to express CD81 fused to mNeonGreen (mNG), enabling fluorescent labeling of secreted EVs. Engineered donor cells were cultured under EV-production conditions, and conditioned media were collected for EV isolation. EVs were purified using size exclusion chromatography (SEC) prior to characterization. Characterization included nanoparticle tracking analysis (NTA) for size distribution, transmission electron microscopy (TEM) for vesicle morphology, imaging flow cytometry for fluorescence and CD81 expression, and Western blot analysis for EV-associated markers. Purified fluorescent EVs were subsequently incubated with different recipient cell lines, including epithelial, immune, and melanoma-derived cells. EV uptake was quantified by flow cytometry based on mNG fluorescence intensity within recipient cells. The figure was created by Biorender.

### 3.1 Characterisation of EVs

To further investigate the potential role of EVs in cellular uptake across different cell lines, EVs were isolated from cultured engineered HEK293T, SKBR-3, and Yummer cells expressing mNG-hCD81 or mNG-mCD81. Vesicles were purified using SEC and subsequently characterized using complementary analytical techniques. Nanoparticle tracking analysis (NTA) demonstrated that the isolated EVs had a modal particle size ranging from 125 to 135 nm, consistent with the expected size distribution of small EVs (Figure 1A, E, F).

Morphological examination by transmission electron microscopy (TEM) confirmed the presence of both small and larger vesicle populations that were predominantly intact, spherical, and membrane bound. Distinct membrane density at the lipid bilayer, together with electron-dense intravesicular cargo, was also visible, further supporting successful EV isolation (Figure 2B, F, J).

**Figure 2.**
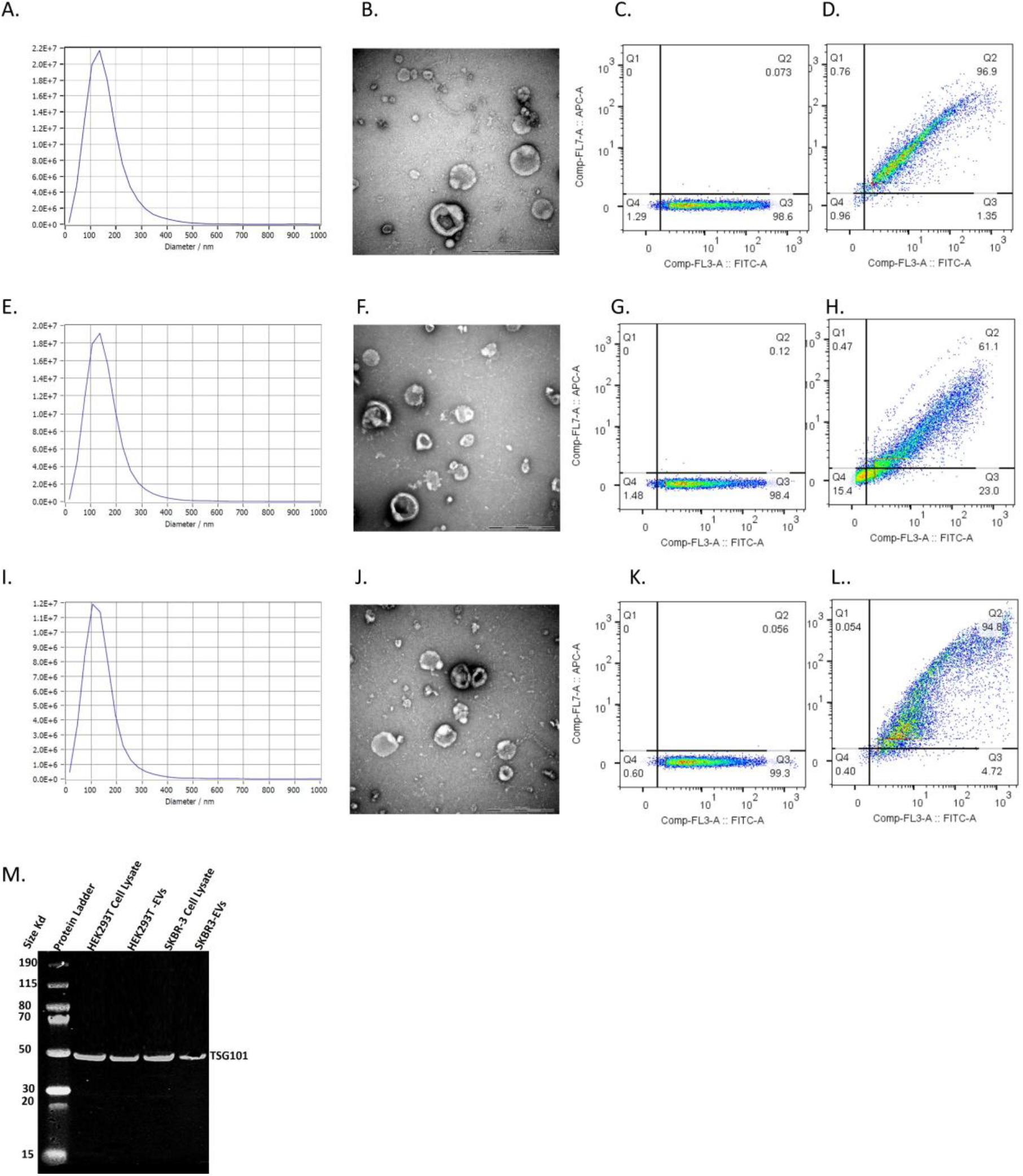
Characterization of extracellular vesicles isolated from engineered donor cell lines. (A) Nanoparticle tracking analysis (NTA) showing the size distribution profile of HEK293T-derived mNG-labeled EVs. (B) Transmission electron microscopy (TEM) image of HEK293T-derived EVs showing intact membrane-bound vesicles. Scale bar = 500 nm. (C) Imaging flow cytometry analysis of HEK293T-derived EVs demonstrating fluorescent mNG-positive vesicles. (D) Imaging flow cytometry analysis of HEK293T-derived EVs stained with APC-conjugated anti-human CD81 antibodies. (E) NTA size distribution profile of SKBR-3-derived EVs. (F) TEM image of SKBR-3-derived EVs. Scale bar = 500 nm. (G) Imaging flow cytometry analysis of SKBR-3-derived EVs. (H) Imaging flow cytometry analysis of SKBR-3-derived EVs stained with APC-conjugated anti-human CD81 antibodies. (I) NTA size distribution profile of Yummer-derived EVs. (J) TEM image of Yummer-derived EVs. Scale bar = 500 nm. (K) Imaging flow cytometry analysis of Yummer-derived EVs. (L) Imaging flow cytometry analysis of Yummer-derived EVs stained with APC-conjugated anti-mouse CD81 antibodies. (M) Western blot analysis of TSG101 in HEK293T-derived and SKBR-3-derived EV preparations.

**Figure 3.**
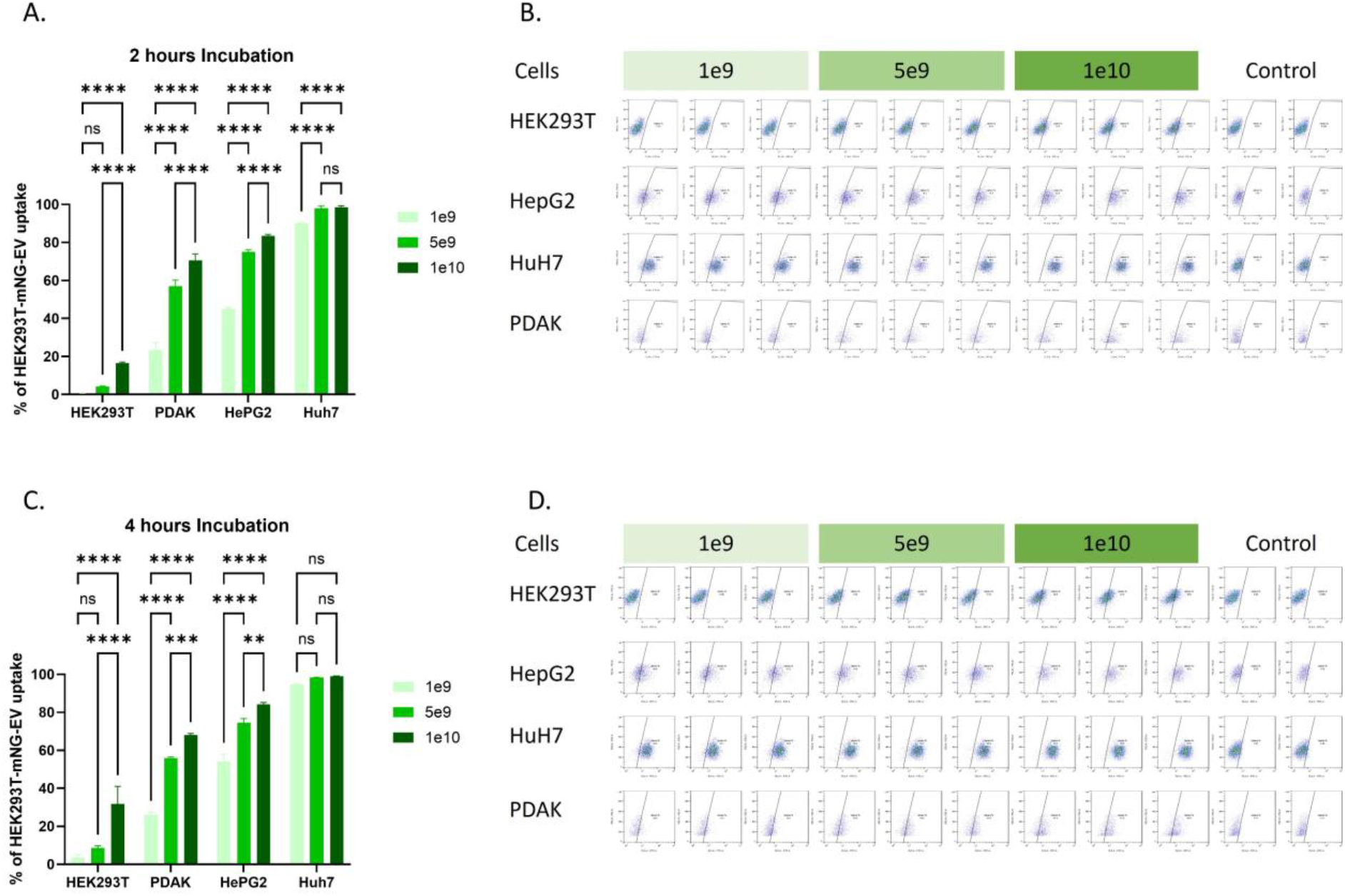
Uptake of HEK293T-derived EVs by human epithelial cell lines. (A) Quantification of EV uptake following 2 h incubation of HEK293T-derived mNG-labeled EVs with epithelial recipient cell lines (HEK293T, PDAK, HepG2, and Huh7) at EV concentrations of 1 × 10^9^, 5 × 10^9^, and 1 × 10^10^ particles. (B) Representative flow cytometry plots corresponding to panel A. (C) Quantification of EV uptake following 4 h incubation at the indicated concentrations. (D) Representative flow cytometry plots corresponding to panel C. Data are presented as mean ± SD (n = 3 independent experiments). Statistical analysis was performed using two-way ANOVA. Statistical significance is indicated as follows: *p < 0.05, **p < 0.01, ***p < 0.001, ****p < 0.0001.

**Figure 4.**
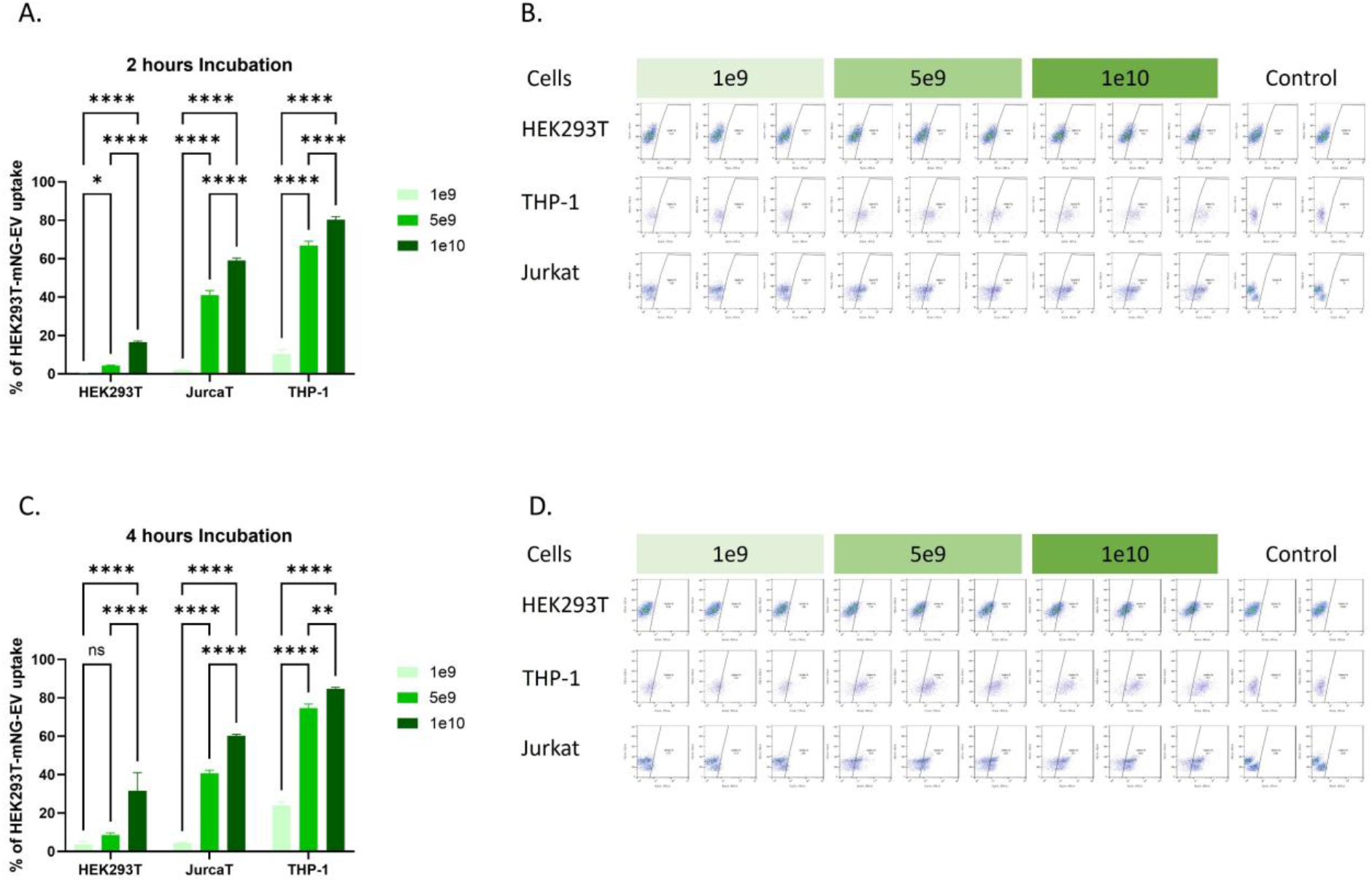
Uptake of HEK293T-derived EVs by human immune cell lines. (A) Quantification of EV uptake following 2 h incubation of HEK293T-derived mNG-labeled EVs with Jurkat, THP-1, and HEK293T cells at EV concentrations of 1 × 10^9^, 5 × 10^9^, and 1 × 10^10^ particles. (B) Representative flow cytometry plots corresponding to panel A. (C) Quantification of EV uptake following 4 h incubation at the indicated concentrations. (D) Representative flow cytometry plots corresponding to panel C. Data are presented as mean ± SD (n = 3 independent experiments). Statistical analysis was performed using two-way ANOVA. Statistical significance is indicated as follows: *p < 0.05, **p < 0.01, ***p < 0.001, ****p < 0.0001.

Flow cytometric analysis revealed that over 94% of the detected vesicles were positive for the canonical EV surface marker CD81, indicating a highly enriched EV population with strong marker expression (Figure 2C-D, G-H, K-L). In addition, Western blot analysis confirmed the presence of tumour susceptibility gene 101 (TSG101), a well-established EV-associated protein, in EV preparations derived from HEK293T and SKBR-3 cells (Figure 2M). Collectively, these findings confirm the successful isolation of structurally intact and marker-positive EVs suitable for subsequent uptake studies.

### 3.2 Uptake of HEK293T-Derived EVs by Human Epithelial Cell Lines

Given that HEK293T-derived EVs are widely used in preclinical applications, it is important to understand whether their internalization differs among recipient cell types. To investigate this, the uptake of HEK293T-derived EVs was evaluated in several human epithelial cell lines in vitro. The results demonstrated that HEK-EV uptake varied substantially between cell lines and depended on both incubation time and EV dose. Experiments were performed at two time points (2 h and 4 h) using three EV concentrations (1 × 10^9^, 5 × 10^9^, and 1 × 10^10^ particles). At the 2 h time point, liver-derived cells exhibited the highest uptake efficiency. Huh7 cells showed the greatest internalization, with uptake values of 90%, 98%, and 98.5% following treatment with 1 × 10^9^, 5 × 10^9^, and 1 × 10^10^ EVs, respectively. HepG2 cells showed the second highest uptake, reaching 45%, 75%, and 85% at the respective doses. PDAK cells ranked third, with uptake values of 23%, 57%, and 71%. The lowest internalization was observed in HEK293T cells, the parental cell line from which the EVs were derived, with uptake values of only 0.6%, 4.5%, and 16.5% at the three tested doses.

At the 4 h time point, EV uptake generally increased. In Huh7 cells, uptake approached saturation levels, reaching 95%, 98.3%, and 99% for the three respective doses. HEK293T cells also showed increased uptake over time, with values of 3.5%, 8.5%, and 31.7%. In contrast, PDAK cells exhibited no notable change between the two time points. Similarly, HepG2 cells maintained uptake values of 75% and 85% at the 5 × 10^9^ and 1 × 10^10^ doses, respectively, whereas uptake at 1 × 10^9^ increased by approximately 10%. Overall, these findings indicate that HEK293T-derived EV internalization is strongly cell type-dependent, with liver-derived cell lines showing the highest uptake efficiency and parental HEK293T cells showing the lowest.

### 3.3 Uptake of HEK293T-Derived EVs by Human Immune Cell Lines

Next, the internalization of HEK293T-derived EVs was investigated in human immune cell lines. THP-1 and Jurkat cells were evaluated in parallel with HEK293T cells as a reference control. Overall, both immune cell lines demonstrated greater EV uptake than HEK293T cells, indicating enhanced internalization capacity of immune cells. At the 2 h time point, THP-1 cells exhibited the highest uptake among the tested immune cell lines, with values of 10.33%, 66.8%, and 80.3% following treatment with 1 × 10^9^, 5 × 10^9^, and 1 × 10^10^ EVs, respectively. Jurkat cells also showed substantial uptake, reaching 2%, 41%, and 59% at the respective doses. In contrast, HEK293T cells displayed the lowest internalization efficiency. At the 4 h time point, THP-1 cells showed a further increase in EV uptake, reaching 23.8%, 74.7%, and 84.7% at the three respective doses. HEK293T cells also demonstrated increased uptake over time, with values of 3.5%, 8.5%, and 31.7% following treatment with 1 × 10^9^, 5 × 10^9^, and 1 × 10^10^ EVs, respectively. However, Jurkat cells showed minimal change compared with the 2 h time point, with only a slight increase (∼2%) observed at the lowest EV dose (1 × 10^9^), while uptake at higher doses remained largely unchanged. Overall, these results indicate that HEK293T-derived EV uptake is more efficient in immune cell lines than in parental HEK293T cells, with THP-1 cells demonstrating the greatest internalization capacity.

### 3.4 Uptake of HEK293T-Derived EVs by Melanoma Cell Lines

As described above, HEK293T-derived EVs were taken up differently by various human cell lines. To determine whether a similar pattern occurs in murine cells, two mouse melanoma cell lines, B16F10 and Yummer, were evaluated in vitro in parallel with HEK293T cells. The results showed clear differences in EV internalization between the tested mouse cell lines. Yummer cells exhibited greater uptake of HEK293T-derived EVs compared with both B16F10 and HEK293T cells. In contrast, B16F10 cells showed lower uptake efficiency than Yummer cells. These findings indicate that EV uptake is strongly influenced by the characteristics of the recipient cell type. This trend was consistently observed at both examined time points (2 h and 4 h) (Figure 5). Furthermore, EV internalization was dose-dependent, as uptake increased with increasing EV concentration. However, once saturation was reached, no significant differences were observed between the higher doses (Figure 5).

**Figure 5.**
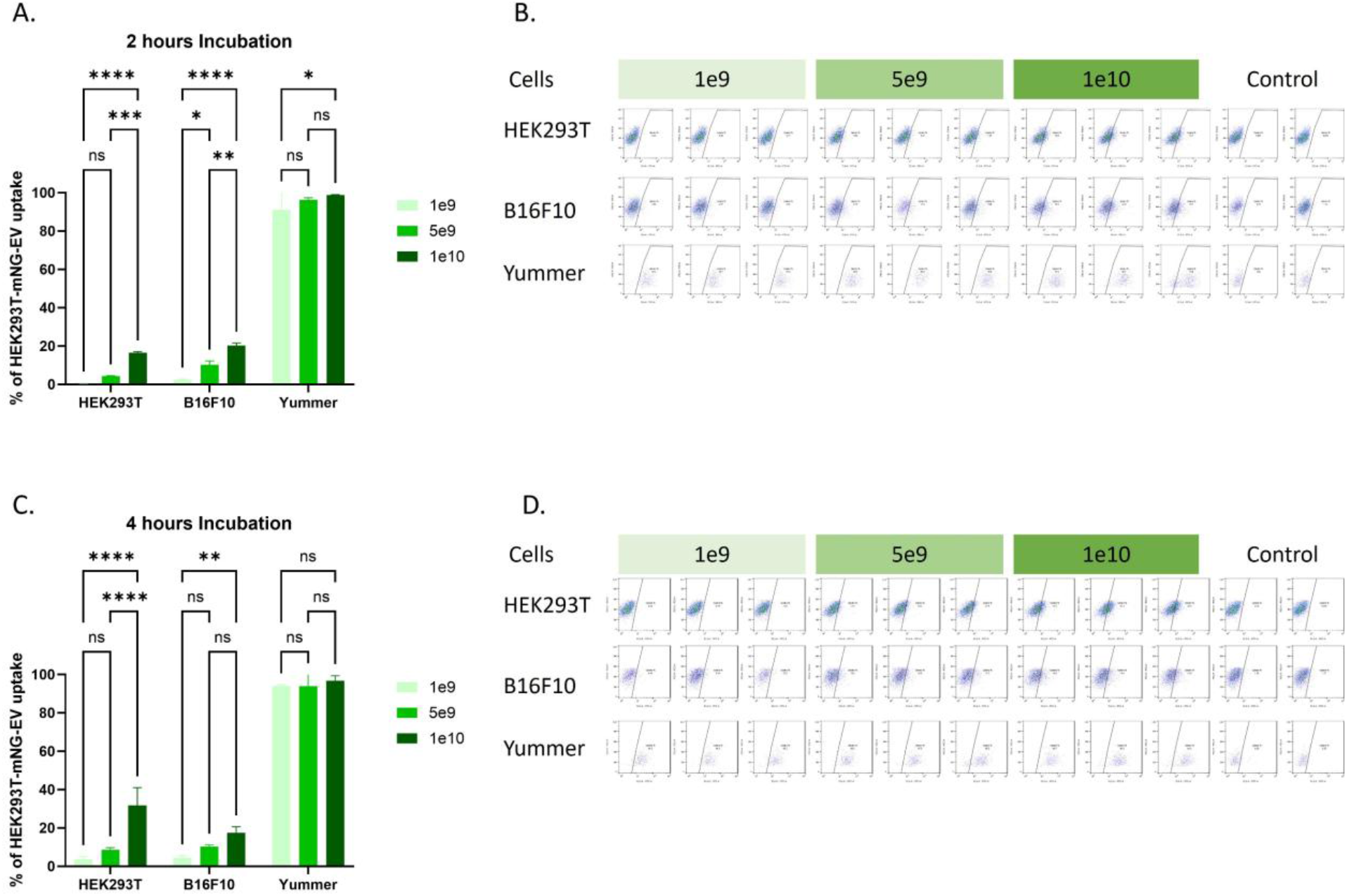
Uptake of HEK293T-derived EVs by murine melanoma cell lines. (A) Quantification of EV uptake following 2 h incubation of HEK293T-derived mNG-labeled EVs with B16F10 and Yummer melanoma cells at EV concentrations of 1 × 10^9^, 5 × 10^9^, and 1 × 10^10^ particles. (B) Representative flow cytometry plots corresponding to panel A. (C) Quantification of EV uptake following 4 h incubation at the indicated concentrations. (D) Representative flow cytometry plots corresponding to panel C. Data are presented as mean ± SD (n = 3 independent experiments). Statistical analysis was performed using two-way ANOVA. Statistical significance is indicated as follows: *p < 0.05, **p < 0.01, ***p < 0.001, ****p < 0.0001.

### 3.5 Uptake of SKBR-3-Derived EVs by Human Epithelial Cell Lines

To investigate the potential role of cellular EV-origin on EV uptake by recipient cells, SKBR-3 cells were transduced with a CD81-mNeonGreen (mNG) lentiviral construct to generate fluorescently labeled mNG-EVs (similar to HEK293T-EVs). These EVs were then incubated with the same panel of human epithelial cell lines (as previously assessed for HEK293T-EVs), including Caco-2, PDAK, HEK293T, SKBR-3, HepG2, and Huh7 cells. The results demonstrated that Huh7 and HepG2 cells exhibited the highest uptake of SKBR-3-derived EVs, exceeding that observed in the EV-producing SKBR-3 cells. PDAK cells also showed increased EV internalization in a dose- and time-dependent manner. A similar pattern was observed in Caco-2 cells, where differences were also evident between EV doses and incubation times. HEK293T cells displayed lower EV uptake compared with most of the tested cell lines. However, they still internalized significantly more EVs than SKBR-3 cells after 4 h of incubation. Overall, these findings further support the conclusion that EV uptake is primarily determined by the characteristics of the recipient cells rather than the cellular source of the EVs (Figure 6).

**Figure 6.**
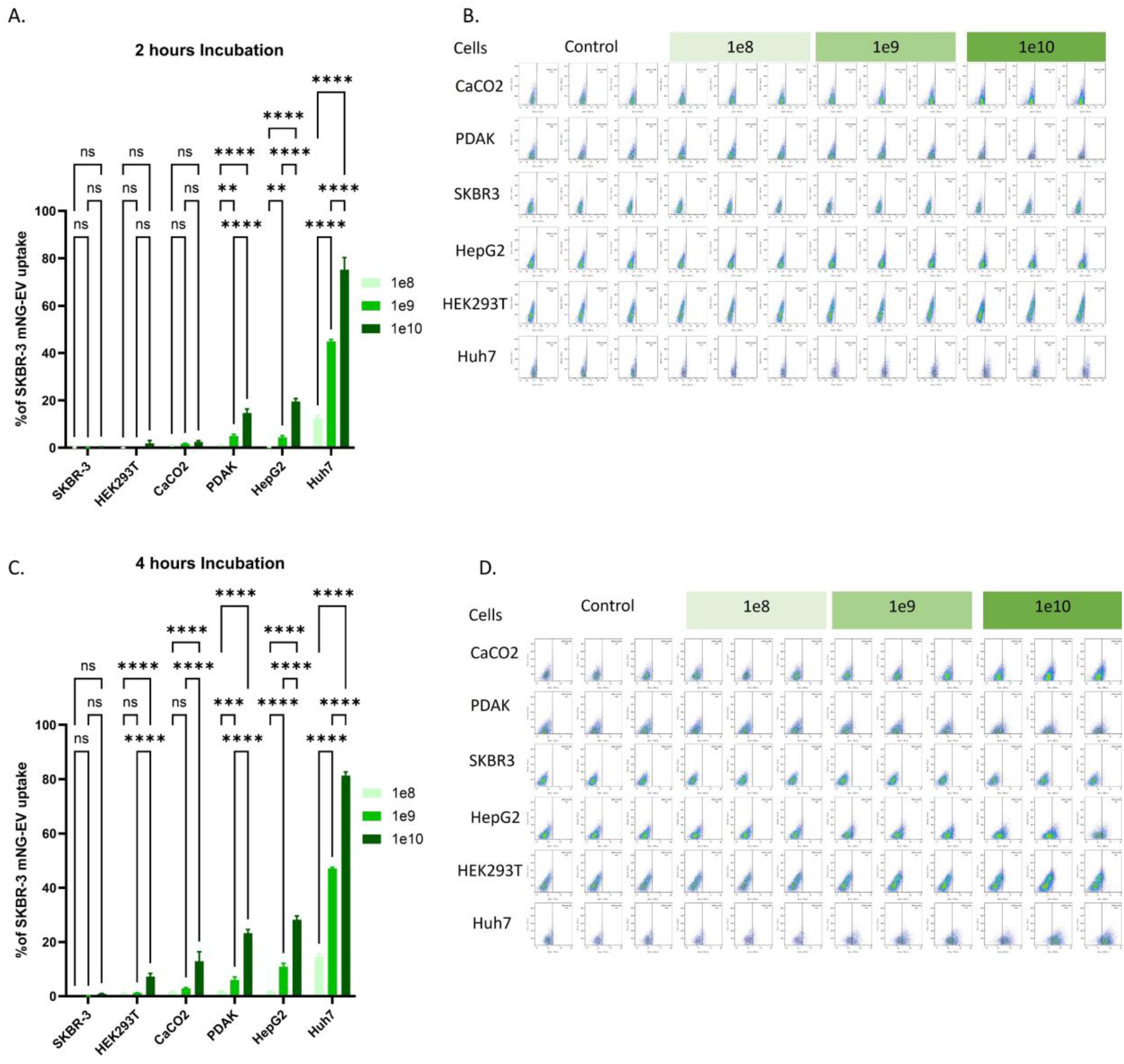
Uptake of SKBR-3-derived EVs by human epithelial cell lines. (A) Quantification of EV uptake following 2 h incubation of SKBR-3-derived mNG-labeled EVs with SKBR-3, HEK293T, Caco-2, PDAK, HepG2, and Huh7 cells at EV concentrations of 1 × 10^8^, 1 × 10^9^, and 1 × 10^10^ particles. (B) Representative flow cytometry plots corresponding to panel A. (C) Quantification of EV uptake following 4 h incubation at the indicated concentrations. (D) Representative flow cytometry plots corresponding to panel C. Data are presented as mean ± SD (n = 3 independent experiments). Statistical analysis was performed using two-way ANOVA. Statistical significance is indicated as follows: **p < 0.01, ****p < 0.0001.

### 3.6 Uptake of Yummer-Derived EVs by melanoma cell lines

Next, we hypothesised that EVs derived from a particular cell source might not be efficiently taken up by the same source or parental cells, based on the above results for HEK293T and SKBR-3 cells, in which EV uptake by the parent cells was low. To test this, Yummer cells were transduced with CD81-mNG lentivirus to generate Yummer-derived mNG-labeled EVs. These EVs were then added to different cell types, including B16F10 and Yummer cells. The results showed that this hypothesis was not universal, as Yummer cells internalized Yummer-derived EVs significantly more than B16F10 cells. These findings confirm that EV uptake is not determined by the cellular source of the EVs but rather depends on the biological membrane properties of the recipient cells (Figure 7).

**Figure 7.**
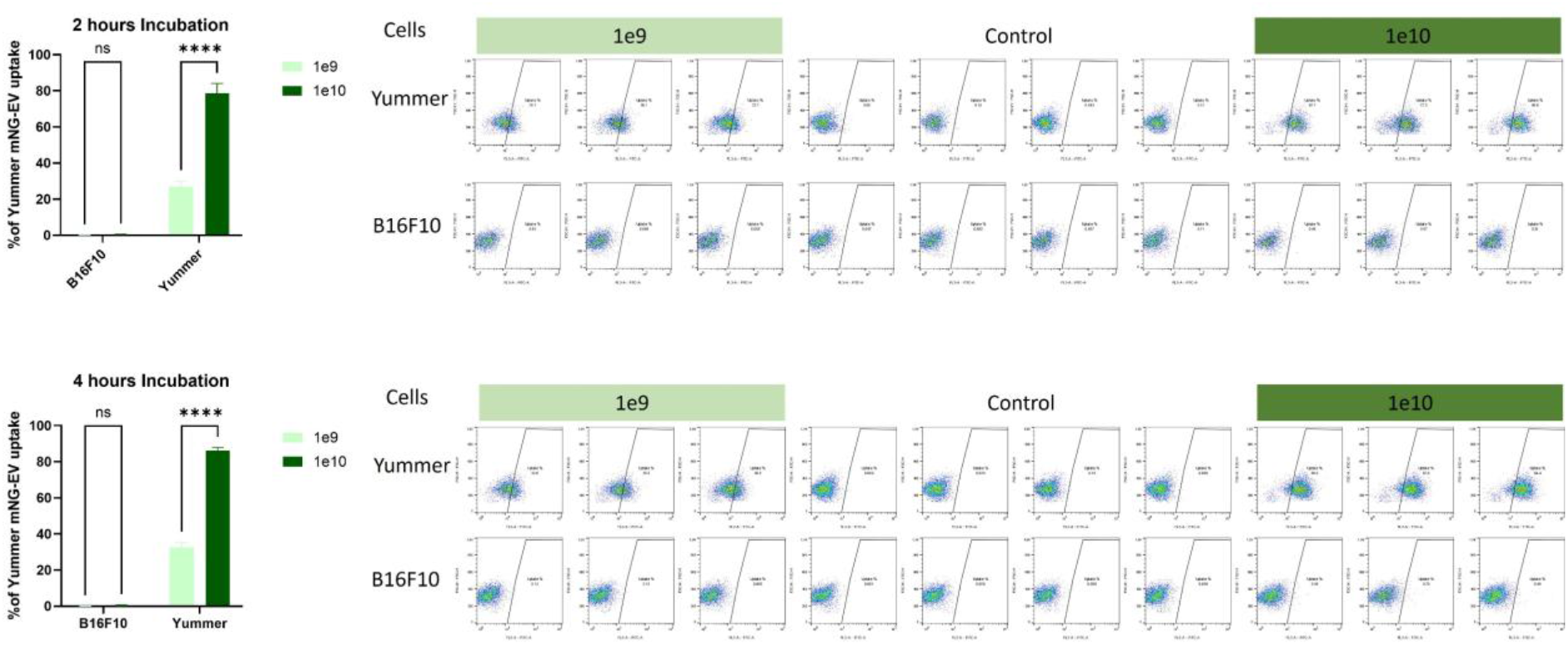
Uptake of Yummer-derived EVs by murine melanoma cell lines. (A) Quantification of EV uptake following 2 h incubation of Yummer-derived mNG-labeled EVs with B16F10 and Yummer cells at EV concentrations of 1 × 10^9^ and 1 × 10^10^ particles. (B) Representative flow cytometry plots corresponding to panel A. (C) Quantification of EV uptake following 4 h incubation at the indicated concentrations. (D) Representative flow cytometry plots corresponding to panel C. Data are presented as mean ± SD (n = 3 independent experiments). Statistical analysis was performed using two-way ANOVA. Statistical significance is indicated as follows: ****p < 0.0001.

## 4. Discussion

EVs are increasingly used as delivery platforms in preclinical and translational research because of their ability to transport bioactive cargo to recipient cells[28]. However, efficient therapeutic application requires a clear understanding of how EVs are internalized by different target cells[29]. In the present study, we systematically compared the uptake of HEK293T-derived and SKBR-3-derived EVs across multiple epithelial, immune, and melanoma cell lines. The data consistently demonstrate that EV internalization is highly dependent on recipient cell type, influenced by dose and incubation time, and only minimally determined by the cellular source of the EVs.

One of the most striking findings was the markedly higher uptake observed in liver-derived epithelial cell lines, particularly Huh7 and HepG2 cells. These cells rapidly internalized HEK293T-derived EVs, reaching near-saturation levels within 2–4 h. This may reflect the naturally high endocytic and metabolic activity of hepatic cells, which are physiologically specialized for uptake of circulating particles, lipoproteins, and macromolecules[30]. The liver is also a major organ for nanoparticle clearance in vivo, and our findings align with previous observations that hepatic cells efficiently internalize nanosized vesicles. This suggests that liver-derived tumors may be particularly responsive to EV-based drug delivery systems, but it also raises the possibility of unwanted hepatic sequestration during systemic administration[31].

In contrast, parental HEK293T cells consistently exhibited the lowest uptake of their own EVs. This is an important and somewhat unexpected observation, as one might assume that cells would efficiently reabsorb vesicles originating from the same cellular background. Instead, the data suggest that EV uptake is not driven by cellular identity matching, but rather by receptor availability, membrane composition, endocytic capacity, and intracellular trafficking pathways in recipient cells. It is possible that HEK293T cells possess fewer surface receptors involved in vesicle docking or have slower membrane turnover compared with more metabolically active cell types[30].

Among immune cells, THP-1 monocyte-like cells showed particularly strong EV uptake, exceeding Jurkat T cells and HEK293T controls. This is biologically plausible, as monocytes/macrophage-lineage cells are professional scavenger cells with robust phagocytic and endocytic machinery. Jurkat cells, although showing moderate uptake, demonstrated less time-dependent increase, suggesting either rapid early saturation or a more limited internalization mechanism[32], [33]. These findings are highly relevant for immunotherapy applications, as EVs intended for tumor targeting may be preferentially captured by circulating immune cells, thereby reducing bioavailability at the intended site.

The murine melanoma experiments further strengthened the concept that recipient-cell phenotype governs uptake. Yummer cells internalized substantially more HEK293T-derived EVs than B16F10 cells, despite both being melanoma-derived lines. This indicates that even closely related tumor models may differ significantly in EV handling. Such differences may arise from variation in membrane lipid composition, expression of heparan sulfate proteoglycans, integrins, tetraspanin-binding partners, or macropinocytosis activity. These data emphasize that tumor type alone cannot predict EV uptake behavior; individual cellular phenotypes must be considered[18].

Importantly, similar uptake patterns were reproduced when the EV source was changed from HEK293T to SKBR-3 cells. Huh7 and HepG2 cells again showed the highest internalization, while SKBR-3 producer cells did not demonstrate superior self-uptake. This cross-validation strongly supports the central conclusion that recipient-cell biology outweighs donor-cell origin in determining EV internalization[18]. Although EV surface proteins and lipid composition likely contribute to biodistribution, their influence appears secondary to the intrinsic uptake capacity of the target cell.

Dose- and time-dependent increases were consistently observed across most cell lines, supporting active uptake mechanisms rather than random surface association. However, saturation occurred at higher EV concentrations, particularly in highly permissive cells. This plateau likely reflects finite receptor availability, limited endosomal processing capacity, or crowding effects at the plasma membrane. From a translational perspective, this is important because escalating EV dose beyond saturation may increase production cost without improving delivery efficiency[34].

Although this study provides a comprehensive comparison of EV uptake across multiple recipient cell types, several limitations should be acknowledged. First, all experiments were performed in vitro, and the observed uptake patterns may not fully reflect EV behaviour in vivo. However, previous in vivo studies have consistently shown that the liver accumulates higher levels of EVs than other organs, supporting our observation that hepatocyte-derived cells exhibit high EV uptake[35]. Similarly, immune cells such as macrophages are also known to efficiently internalize EVs in vivo[36]. Second, the molecular mechanisms underlying differential EV uptake were not directly investigated, and no inhibition studies were performed to identify specific internalization pathways. Future studies should therefore focus on mechanistic investigations and in vivo validation to further support the translational potential of EV-based therapeutics.

Collectively, these findings have direct implications for EV-based therapeutics. First, recipient cell selection is a major determinant of delivery success. Second, biodistribution may be heavily biased toward highly endocytic tissues such as liver and immune compartments. Third, engineering EV donor cells alone may be insufficient unless recipient-cell targeting mechanisms are also optimized. Future studies should therefore focus on defining the molecular determinants of selective uptake and developing EV surface modifications that overcome natural sequestration patterns.

## 5. Conclusion

This study demonstrates unequivocally that extracellular vesicle uptake is governed primarily by the biological properties of recipient cells rather than by the cellular source of the vesicles. Across human epithelial, immune, and murine melanoma models, EV internalization varied markedly between cell types, while similar uptake hierarchies were maintained even when the EV donor cell line was changed. Liver-derived cells and monocyte-like immune cells showed the greatest uptake efficiency, whereas parental donor cells often displayed poor self-internalization. Uptake was further enhanced by increasing dose and incubation time until saturation was reached. These findings establish recipient-cell phenotype as the dominant factor controlling EV delivery and highlight a critical consideration for the rational design of EV-based therapeutics: successful targeting will depend less on where EVs come from and more on where they are going.

## Acknowledgements

I would like to thank Oscar Wiklander and his group for their support of the research described in this study. This work was supported by the Swedish Research Council (VR, 2022-02449), the Swedish Cancer Society (project 23 2935 Pj), Radiumhemmet Research Funds (project #241392), the Center for Innovative Medicine junior investigator grants (FoUI-976434), and Karolinska Institutet (2-116/2023). I would also like to thank the Division of Biomolecular and Cellular Medicine for providing the facilities that made this study possible and for allowing me to use the facility. I would also like to acknowledge Microsoft Copilot and Open AI ChatGPT, which were used solely for proofreading and language editing purposes. The author has reviewed the entire text and takes full responsibility for the content of the manuscript. The author would also like to acknowledge Dr. Oscar Wiklander for his valuable review and editing of the manuscript.

## Author contributions

Doste R. Mamand: Conceptualization, data curation, formal analysis, investigation, methodology, resources, supervision, visualization, writing—original draft, writing—review, and editing.

## Data Availability Statement

The data that support the findings of this study are available from the corresponding author upon reasonable request.

